# Reversible kink instability drives ultrafast jumping in nematodes and soft robots

**DOI:** 10.1101/2024.06.07.598012

**Authors:** Sunny Kumar, Ishant Tiwari, Victor M. Ortega-Jimenez, Adler R. Dillman, Dongjing He, Yuhang Hu, M. Saad Bhamla

**Affiliations:** School of Chemical and Biomolecular Engineering, Georgia Institute of Technology, Atlanta, GA 30332; Department of Integrative Biology, University of California, Berkeley, CA 94720; Department of Nematology, University of California, Riverside, CA 92521; Department of Mechanical Engineering, Georgia Institute of Technology, Atlanta, GA 30332

## Abstract

Entomopathogenic nematodes (EPNs) exhibit a bending-elastic instability, or kink, before becoming airborne, a feature hypothesized but not proven to enhance jumping performance. Here, we provide the evidence that this kink is crucial for improving launch performance. We demonstrate that EPNs actively modulate their aspect ratio, forming a liquid-latched closed loop over a slow timescale *O*(1 s), then rapidly open it *O* (10 µs), achieving heights of 20 body lengths (BL) and generating ∼ 10^4^ W/Kg of power. Using jumping nematodes, a bio-inspired Soft Jumping Model (SoftJM), and computational simulations, we explore the mechanisms and implications of this kink. EPNs control their takeoff direction by adjusting their head position and center of mass, a mechanism verified through phase maps of jump directions in simulations and SoftJM experiments. Our findings reveal that the reversible kink instability at the point of highest curvature on the ventral side enhances energy storage using the nematode’s limited muscular force. We investigated the impact of aspect ratio on kink instability and jumping performance using SoftJM, and quantified EPN cuticle stiffness with AFM, comparing it with *C. elegans*. This led to a stiffness-modified SoftJM design with a carbon fiber backbone, achieving jumps of ∼25 BL. Our study reveals how harnessing kink instabilities, a typical failure mode, enables bidirectional jumps in soft robots on complex substrates like sand, offering a novel approach for designing limbless robots for controlled jumping, locomotion, and even planetary exploration.

## A Cylindrical Shape Motif Forms Life

The cylindrical shape is a hallmark of biological systems, evident across length scales and taxa, from bacteria to worms to plants and animals (including humans), and their appendages, stems, tails, antennae, and flagella [1]. Kinks, a universal phenomenon, occur when the curvature of a cylindrical structure’s compressed side exceeds a critical value, leading to localized instability. This can be illustrated by bending a plastic straw, where the localized curvature exceeds a critical point, causing a kink [2, 3]. In biological systems, kinks are usually detrimental: they can cause permanent damage in plant stems [4], blockages in blood vessels [5], and failures in insect exoskeletons [6]. Despite their typically harmful effects, in this work, we report a functional use of kink instability by a living system —a nematode—for high-powered aerial jumping locomotion (Fig. 1A-C).

**FIG. 1.**
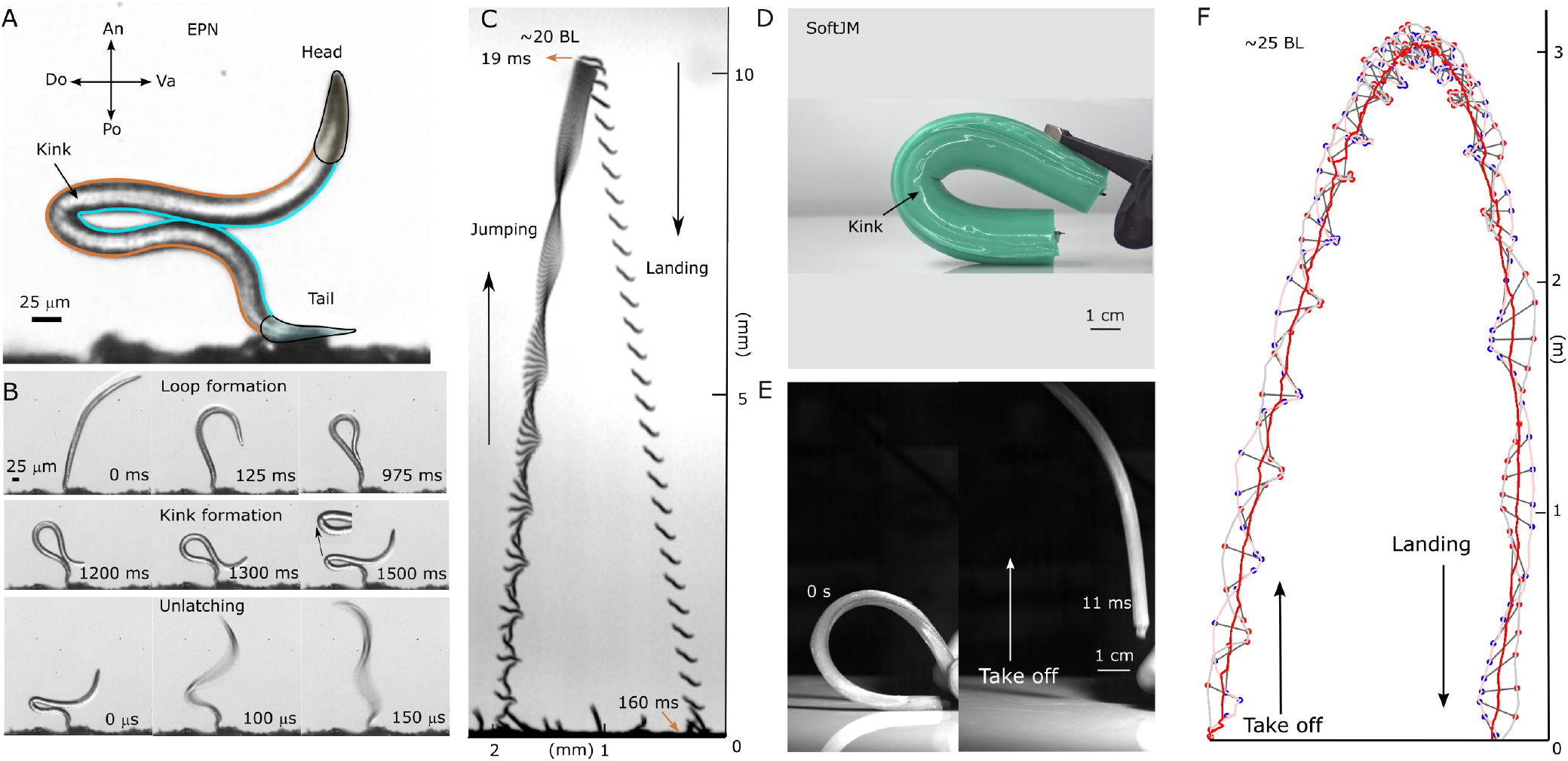
Jumping Nematodes and Bio-inspired Soft Jumping Model. (**A**) Image of a *Steinernema carpocapsae* nematode in a bent or “kink” position, captured just before jumping, as seen through a high-speed camera with a zoomed-in view. The scale bar is 25 *µ*m (Po-Posterior, Do-Dorsal, An-Anterior, Ve-Ventral). (**B**) Sequential images illustrating the EPN’s jump preparation: loop formation from 0 to 975 ms, kink formation from 1200 ms up to 1500 ms, and final unlatching phase lasting 150 *µ*s. The scale bar is 25 *µ*m. (**C**) Trajectories of the EPN during take-off (speed ∼1.5 *m/s*), rotation, and landing (speed ∼0.15 m/s). (**D**) Soft jumping model (SoftJM) representing a physical model to replicate the EPN’s jump at the macro scale. (**E**) Images illustrating the takeoff phase of the SoftJM. (**F**) Trajectory of the SoftJM showing takeoff, rotation, and landing phases.

### Harnessing Elastic Instabilities in Soft Robotics

Elastic instabilities, traditionally seen as failure modes, are now recognized as useful mechanisms in the transition from rigid to more compliant structures[7]. In soft actuators, these instabilities have enabled the creation of functional devices such as snap-through inflatable shell actuators [8], and kink-valves in pneumatic tubes[9]. However, kink instability, most commonly observed in cylindrical rods and underpinning other classes of instabilities such as the Brazier instability, remains largely underutilized in jumping soft robots. Here, we demonstrate a nematode-inspired soft cylindrical limbless jumper (Fig. 1D-F) that harnesses kink instabilities to achieve effective jumping performance.

### Microsecond Kink Instability in Tiny Nematodes

Nearly 50 years ago, researchers documented the remarkable jumping ability of the limbless, soft-bodied EPNs [10]. Subsequent observations also noted a kink in its body [11, 12]. However, the functional role of this kink remained unexplored and under-appreciated due to the lack of high-speed microscopy, since the kink-opening timescales are in microseconds. To address this gap, we cultured EPNs and conducted high-speed experiments, revealing the role of the kink in unprecedented detail.

Our observations focused on the infective juvenile (IJ) stage of *Steinernema carpocapsae*, which is about 20 *µ*m wide and 0.5 mm long, yielding a highly anisotropic slender object of aspect ratio of 25-30. Under the right conditions of humidity and olfactory stimulus [13–15], the nematode can stand up and contort its body into a ventral-ventral contact, forming an *α*-shape. Using a high-speed camera, we observed that the EPN can tighten the ventral-ventral loop in approximately 1.5 seconds by moving its head forward, leading to the formation of a kink in its compressed (ventral) side. Subsequently, it opens the loop in about 150 *µ*s (about 50 times faster than the snap of a finger [16], actuating a jump *>* 20 BL high, with a take-off speed of approximately 1.5 m/s (Reynolds number Re∼22, Fig. 1C, Movie S1, Fig. S2). Initially, the EPN rotates rapidly (about 2000 rotations/s), slowing down to almost zero rotation as it reaches the apex of the jump, and then falls slowly with a speed of 0.15 m/s (Re∼1, SI Methods and Table S6) [17]. The ballistic jump of the nematode follows an asymmetric trajectory, resembling the Tartaglia curve, where the take-off velocity is greater than the falling velocity[18]. This trajectory allows the nematode to maximize its chances of reaching a host without overshooting it horizontally.

### Directional Control in EPN Jumps

Can these effectively limbless, cylindrical ambush-forager EPNs control the direction of their jump? We discovered that the EPN can take off in forward, upward, and even backward directions. Two stereotypical examples of this jump are presented in Fig.2A (left panel) and Movie S1. The EPN shifts its center of mass (COM) by adjusting the orientation of its head *α* and its loop *θ* to bias the take-off angle *γ* either forward or backward [19]. This reveals an elegant and simple principle for any elastic cylindrical structure to control the direction of its jump. To demonstrate the generality of this posture-based COM-directional control in elastic filaments, we validated this COM-controlled jump direction via in-silico simulations of a Cosserat rod [20, 21] at the micron scale and a physical SoftJM at the macro scale. The EPN and the Cosserat rod have similar maximum angular opening speeds 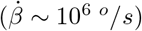 due to their micron length scale, while the SoftJM opens relatively slowly 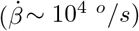 since it actuates at the centimeter scale (Fig. 2B).

**FIG. 2.**
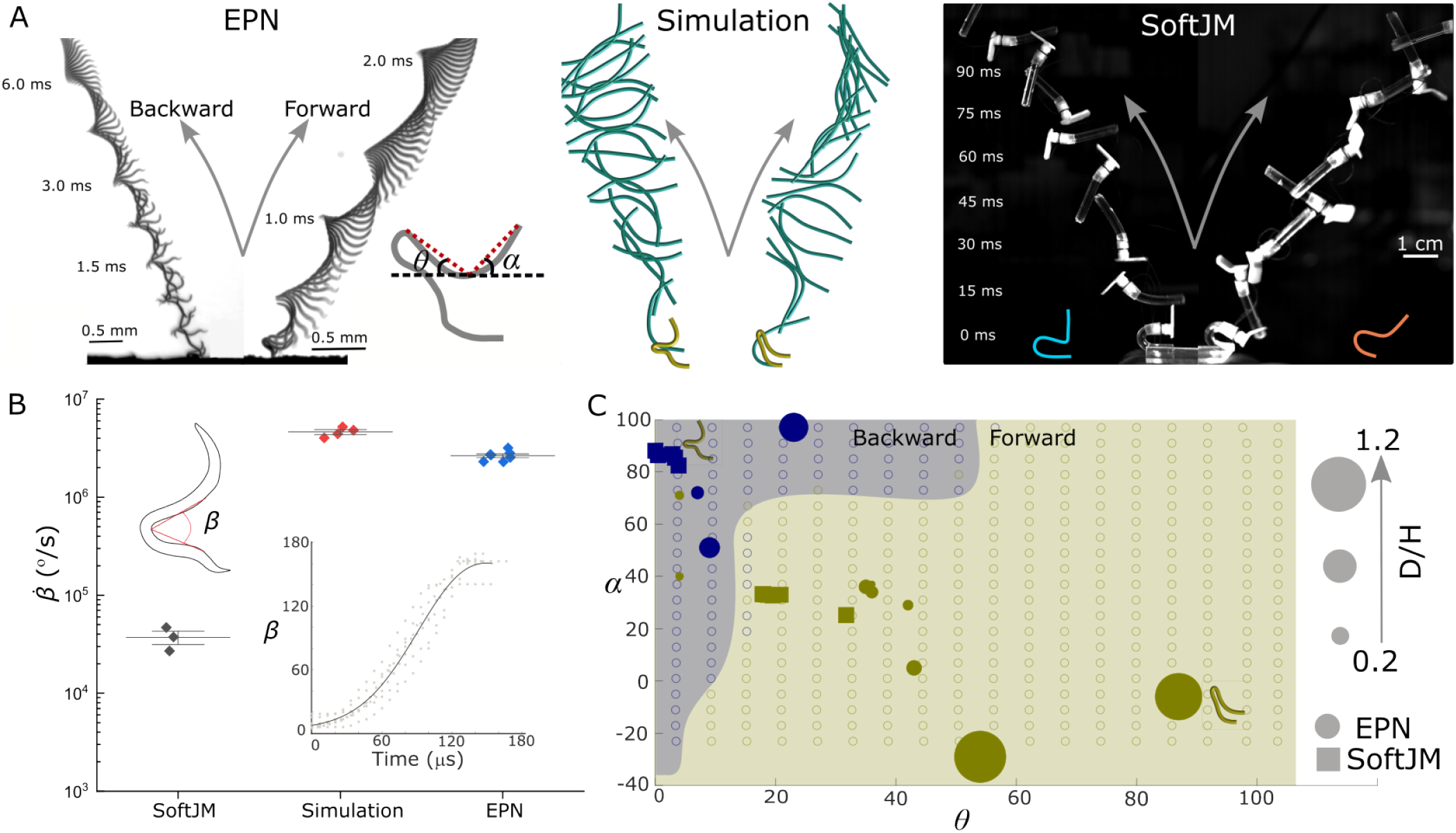
Directional Jumping of the EPN, SoftJM and Simulation. (**A**) Jumping of the EPN in the backward and forward directions. The EPN’s take-off angle depends on its posture, quantified by the angles *θ* and *α*. Adjusting these angles can move its center of mass backward or forward with respect to its point of contact with the substrate, resulting in a backward jump (*α* ∼ 90^*°*^, *θ* ∼ 0^*°*^, and *γ* ∼ 120^*°*^) or forward jump (*α* ∼ 45^*°*^, *θ* ∼ 45^*°*^, and *γ* ∼ 60^*°*^) with additional details in Table S2 and S5. Cosserat rod simulations and physical models of soft jumpers also show bidirectional jumping. (**B**) Angular velocity (*β*?) for SoftJM (n=3 individuals), simulation (n=4 realizations), and EPN (n=7 individuals). The inset shows the opening angle *β* plotted as a function of time for the EPN (n=5). (**C**) Simulation results of different postures of the simulated EPN with different color representations for backward (blue rings) and forward (yellow rings) take-offs. Solid blue (yellow) color depicts experimental jumps in the backward (forward) direction, with circles representing EPN jumps while squares represent SoftJM jumps. The size of these solid-colored data points is proportional to their normalized trajectory range (D/H) (see details in Fig. S2 and S3).

We explored a multitude of initial jump postures beyond the capabilities of the EPN by creating a numerical phase space of jump directions, controlling the location of the COM using the head angle *α* and the loop angle *θ* (Fig. 2C). The phase space is divided into forward and backward jumping regions, separated by postures that jump vertically. When measuring the head and the loop angles of the EPN and the SoftJM and placing them on this map, we found that all the experimental points lie on a line, indicating that experimentally, there are constraints coupling the head and the loop angles. This finding implies the possibility of designing novel jumping robots that can exploit the available phase space more effectively (see Discussion).

### Stiff Cuticle and *α*-Shape Spring-Loads EPN Jumps

What structural and mechanical properties enable parasitic EPNs to jump, unlike similar sized nematodes like *C. elegans*? Small-scale organisms often exhibit springlatch actuation to perform activities at power levels significantly higher than their muscular capabilities [23, 24]. Notably, we found that the maximum speeds observed during the jump of an EPN are more than 100x larger than those observed during nictation and loop formation (Fig.3D, Movie S2), indicating that the jumping is driven by a spring-latch mechanism rather than purely muscle power (∼ 10^4^ W*/*Kg, details in SI materials and method sections). The IJ (dauer) stage of the EPN has a thick cuticle enclosing a hydrostatic skeleton that could act as a spring to store energy during the jump [25]. To confirm this, we conducted AFM measurements on dauer stages of both EPN and *C. elegans*, revealing a ∼10x higher bending stiffness in EPNs (Fig. 3E and Fig. S1). This stiffer (and thicker) cuticular structure likely enables *S. carpocapsae* to jump, unlike other nematodes with similar structural properties, such as *C. elegans*.

**FIG. 3.**
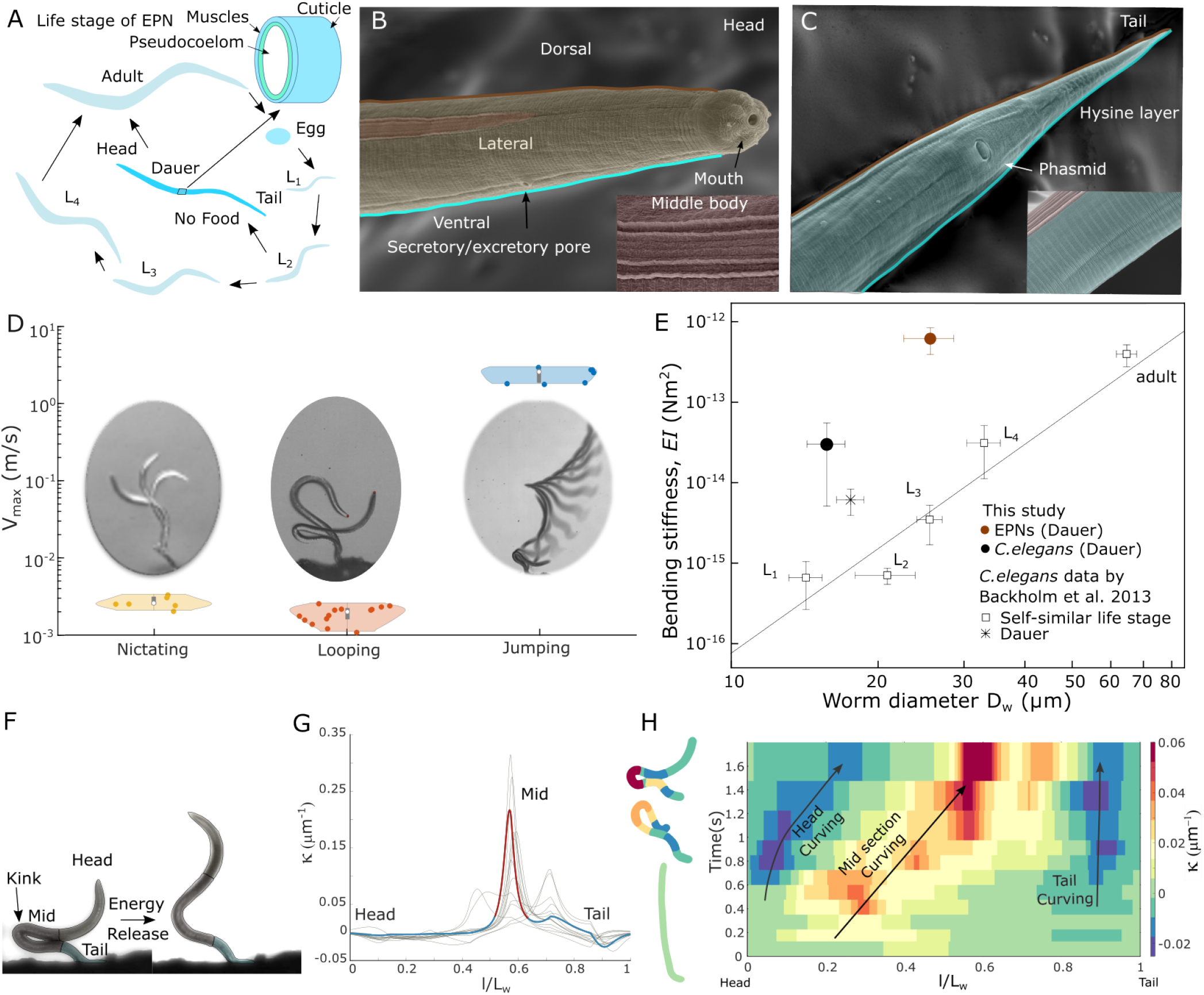
Bending Stiffness and Curvature Analysis of Nematodes. (**A**) Life stages of the EPN from L1, L2, L3, L4, dauer stage (IJ), and adult stage. The cross-sectional schematic of the dauer stage nematode shows the complex arrangements of the cuticle, muscles, and pseudocoelom. (**B**) Scanning electron microscopy (SEM) of EPN, highlighting the head and neck region of infective juveniles (dauer). The yellow arrow marks the mouth and secretory pore in the head region. The inset shows the lateral side of the EPN body. (**C**) SEM of the EPN’s tail, with the inset showing the ventral side of the nematode. (**D**) Maximum speed distribution of the EPN during nictating (n=6), looping (n=14), and jumping(n=6). (**E**) Bending stiffness of the dauer stage EPN and *C. elegans*, measured by atomic force microscopy (AFM) as a function of nematode diameter *D*_*w*_ (n=3 individuals, 3 times each). The comparison includes self-similar nematodes averaged over each life stage for *C. elegans* reported by Backholm et al. 2013 [22]. The dauer stage EPN shows higher stiffness compared to the same self-similar life stage and dauer of *C. elegans*. (**F**) Different regions (head, middle, and tail) of the EPN before jumping and opening the loop. (**G**) EPN’s final state curvature with normalized length (l/L_*w*_) before jumping (n=14), where *l* is the varying length across the nematode body, and L_*w*_ is the total length of EPNs. (**H**) The kymograph shows the curvature growth across the length of the EPN as a function of time. The color-coded EPN schematic on the left visualizes the curvature profile along the body of the nematode.

To investigate how EPN harnesses its body as a spring to store energy, we quantified the geometry of the EPN during the jump. This analysis helped identify where the nematode stores energy for the jump. Right before its jump, the EPN’s geometry shows maximum curvature in the middle of its body, where the kink forms (Fig. 3F, G). As the nematode forms the loop, only the mid-section accumulates curvature while the head and the tail region remain relatively unchanged (Fig. 3H). Post-jump, the mid-section curvature is released, while the head and tail remain curved. These findings suggest that EPNs leverage both a stiff cuticle and their *α*-shape geometry as a spring.

### Liquid Latch Facilitates EPN’s *µ*s Release

For this microscopic nematode (thinner than a human hair), what kind of ‘latch’ holds this *α*-shape spring to-gether for a quick microsecond release? Latches, despite their ubiquity in man-made systems, have been relatively understudied in biology [16, 23, 26]. In the case of EPNs, it has been hypothesized (but not proven) that a water droplet acts as a liquid latch to hold the ventral-ventral contact of the EPNs *α*-shaped spring [10] (Fig. 4A). Using high-speed video recordings of the nematode, we observe, for the first time, the rupture of a liquid latch as the EPN opens its loop (∼ 70*µs*, Fig. 4D, Movie S2). We create a theoretical model of the capillary force [27] (*F*_*latch*_, Fig. 4B,C) that can balance the force generated from the *α*-shaped curvature [28, 29] of the EPN (*F*_*curve*_)

**FIG. 4.**
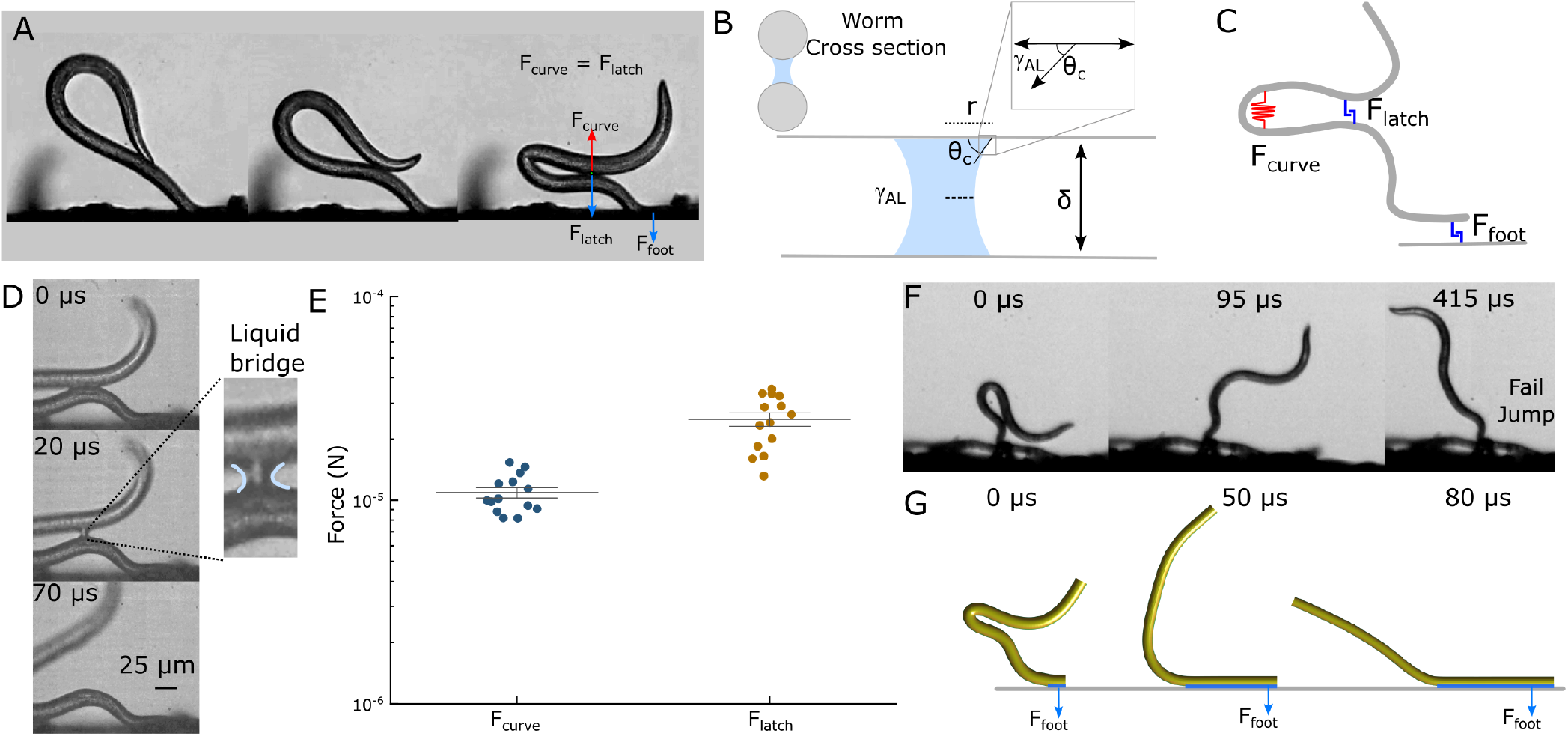
Liquid Latch Mechanism and Force Balance. (**A**) EPN forming the liquid latch at ventral-ventral contact. (**B**) Cross-sectional image of the liquid bridge where the surface tension of the liquid is denoted by *γ*_*AL*_, assuming perfect wetting (*θ*_*c*_ = 0). (**C**) A simplified schematic showing the forces involved in the EPN’s jump: capillary latch *F*_*latch*_, curvature force *F*_*curve*_, and tail adhesion due to capillary forces *F*_*Foot*_. (**D**) Liquid bridge thinning and breaking over time during ventral-ventral contact of EPN. (**E**) Bending force *F*_*curve*_ due to curvature and capillary force *F*_*latch*_ due to the liquid bridge between the ventral-ventral contact (n=15), details in Table S7. (**F**) Failed jump due to tail adhesion to the surface. (**G**) Simulation snapshots illustrating the failed jump caused by simulated adhesion on the tail.

defined as:

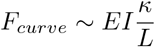

and

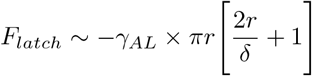

where *EI* and *κ* are the bending stiffness and the curvature of the EPN, respectively, while *γ*_*AL*_ is the surface tension of the air-water interface, 2*r* is the ventral-ventral contact length, and *δ* is the height of the capillary-latch right before the jump (See Fig. 4B and detailed derivation in Supplementary text). Using the parameters for the worm (Table S7), we found that the latch force and maximum curvature force are of the same order of magnitude (*F*_*curve*_ = *F*_*latch*_ ∼ 10^−5^ N). This reinforces the idea that in this nematode, a few microns thick liquid film can act as a sufficient latch to hold the curved ends of the worm in place until ready to take off. The EPN remains in the *α*-loop position until the force magnitudes balance each other (Fig. 4E). The jump occurs when *F*_*curve*_ *> F*_*latch*_, breaking the latch and rapidly opening the nematode’s loop in ∼ 70*µs* microseconds. It is worth noting that surface tension has been exploited as a ‘spring’ and ‘engine’ by other invertebrates [30, 31], and our work adds to this by revealing how EPNs use surface tension as a functional ‘latch’.

Another liquid latch is present between the tail of the EPN and the substrate. If the force due to this liquid bridge is high enough, the EPN can fail to leave the substrate even with the release of the liquid latch at the ventral-ventral contact point (Fig. 4F). Similar observations are made in our computational model by introducing an attraction between the foot of the Cosserat rod and the substrate (Fig. 4G). Our experimental observations and its relatively simple numerical reproduction indicate that if the humidity is too high and a large liquid bridge forms between the tail and the wet substrate, the EPN can fail its jump even after a successful *α*-loop opening (Fig. S4 and Movie S2).

### Soft JMs Store More Energy Via Reversible Kinks

How does the kink instability help the EPN store energy for its jump? We answer this question using scaled physical models (SoftJMs) because similar measurements are inaccessible at the micron scale of the EPN. We test 4 types of synthetic SoftJMs (Fig. 5A): a hydrostatic skeleton body (SoftJM 1), a hydrostatic skeleton with a stiff backbone (SoftJM 2), a soft PDMS rod (SoftJM 3), and a soft silicone rod with a stiff backbone (SoftJM 4).

**FIG. 5.**
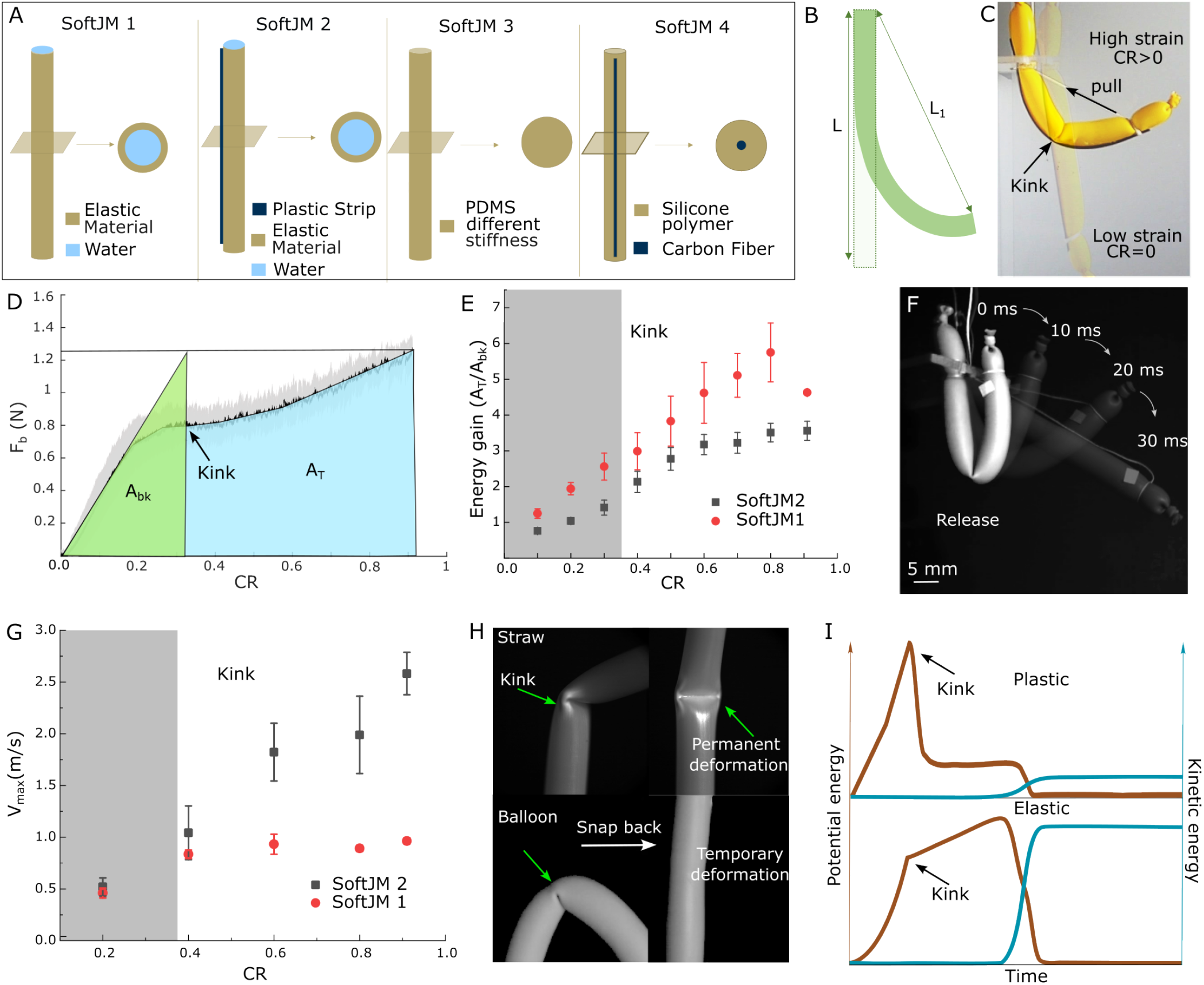
Kink-Induced Elastic Energy Storage and Force Dynamics in SoftJMs. (**A**) Various synthetic soft jumping models (SoftJM): SoftJM 1 (Water Balloon, low stiffness), SoftJM 2 (Water balloon + Strip, stiff backbone), SoftJM 3 (PDMS, different Young’s modulus, see Table S4), SoftJM 4 (Silicone Polymer + Carbon Fiber (CF)). (**B**) Schematic for bending of rod to measure bending force (*F*_*b*_), where Compression Ratio, *CR* = (*L* − *L*1)*/L*. (**C**) Kink formation during bending of SoftJM 2, aspect ratio *η* = 10. (**D**) Bending force *F*_*b*_ with different compression ratios (CR) for SoftJM 2 (black line) *η*=10. (**E**) Energy gain (*A*_*T*_ */A*_*bk*_) for both SoftJM 1 and SoftJM 2 (n=5), *η*=10. (**F**) Snap back visualization of SoftJM 1 at different time intervals, *η*=10. (**G**) Maximum velocity (*V*_*max*_) graph illustrating the maximum snapback velocity for different physical models, SoftJM 1 (red points) and SoftJM 2 (black points) (n=3), *η*=10. (**H**) A cylindrical rod bends and snaps back if there is temporary deformation (elastic balloon or PDMS); otherwise, it undergoes permanent deformation (plastic straw). (**I**) Schematic illustration of energy balance representing input potential energy (red line) and output kinetic energy (blue line) for irreversible (plastic) and reversible (elastic) systems.

Using a custom-built rig, we bend SoftJM 2 and measure its restoring force as a function of the bending strain, quantified by the compression ratio (CR). The restoring force grew linearly until a kink formed, beyond which the force continued to increase, albeit at a lower slope (Fig. 5B-D, Movie S3). This nonlinear response to bending strain allows the storage of excess energy in systems limited by a force ceiling, which is common in biological motors [32]. This energy gain can be estimated assuming a force ceiling as the maximum force observed in an experiment and then extrapolating the initial linear curve in restoring force (pre-kink) to this force ceiling. The energy gain is effectively a comparison of the area under the curve for the actual force curve and the linear extrapolated curve, which is ∼ 5x and ∼ 4x for SoftJM 1 and SoftJM 2 (Fig. 5E), compared to the hypothetical linear spring scenario of no kink formation. Similar force measurements for PDMS (SoftJM 3) of varied stiffness coefficients and stiff plastic strips are presented in Fig. S5.

Simply reducing force by kinking in an elastic structure is insufficient. Power generation during actuation is equally critical for a successful jump. We thus measure the maximum opening speed V_*max*_ of the SoftJM’s as a metric of the output power. V_*max*_ almost doubled when the bending strain increased from CR=0.4 to CR=0.9, indicating that the power output of the jump increases even after the formation of the kink (Fig. 5G).

Not all kink instabilities benefit actuation. If a plastic material undergoes *irreversible* deformation post-kink, such as in a drinking straw, undersea pipelines, or rodshaped *E. coli* bacteria [3], the kinetic energy released during relaxation is minimal (Fig. 5H,I). The EPN and the SoftJM show a *reversible* kink formation, enhancing their elastic energy storage and subsequent release of kinetic energy (Fig. 5I). Therefore, the occurrence of reversible kink instability in the EPN may help it store a larger amount of elastic energy while staying within a finite budget of muscle force.

### EPNs Actively Modulate Aspect Ratio and Its Effect on SoftJMs

The aspect ratio, *η* = *L/D*, defined as the length-to-diameter ratio, is critical for the stability of cylindrical structures. Cylinders with high *η* are more susceptible to buckling instability, while those with low *η* are harder to bend, typically requiring more force. We discover that EPNs actively modulate their aspect ratio via their *α*-loop, starting from an initial *η* ∼ 20 to *η* ≲ 7, reducing their loop aspect ratio by nearly 3x (Fig 6A,B). By reducing their effective *η* dynamically, the EPNs achieve a two-fold benefit: they increase their midsection curvature (*κ*) by 2-3 times, storing elastic energy in their bent bodies at high forces; and upon reaching the upper force-limit of the bending near the critical *η* ≲ 7 and *κ* ≥ 0.025*µ*m^−1^, they form kinks, allowing them to continue loading energy without an increase in force. This process enables EPNs to ‘charge’ up more elastic energy in the bent configuration as they prepare to jump.

**FIG. 6.**
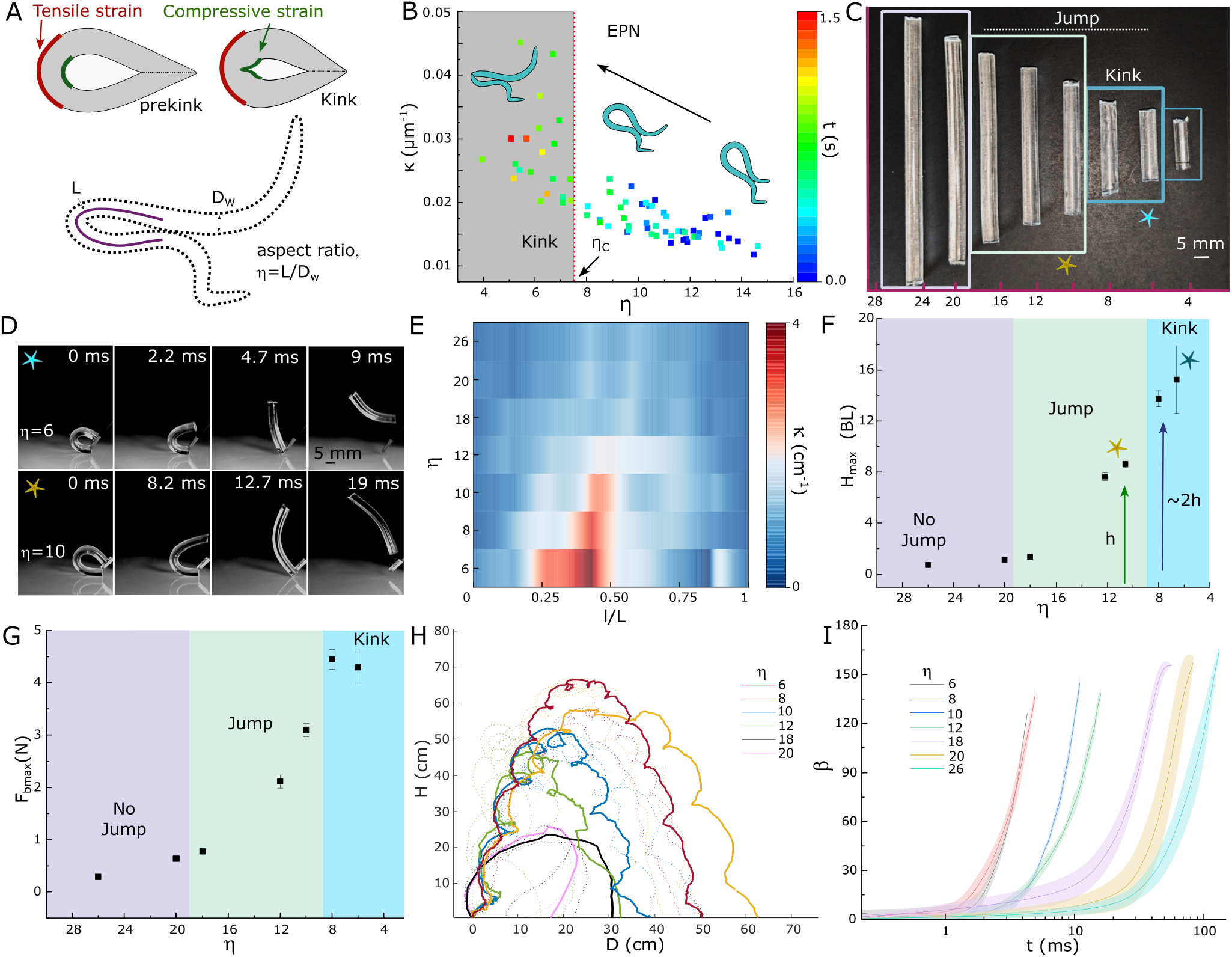
Role of Aspect Ratio in EPN and Soft JM. (**A**) EPNs’ geometrical configuration and transition from closed loop to kink formation, where the aspect ratio is defined as *η* = *L/D*_*w*_ . During bending or compressive stress, the outer surface of the EPN experiences tensile strain while the inner surface experiences compressive strain. At a critical degree of bending, a sharp kink appears on the inner surface. (**B**) EPNs’ geometrical transition from loop to kink, representing the relationship between curvature *κ* and *η* (n= 8 individuals). (**C**) SoftJM 3 at different *η* for modulus of 1.2 MPa. (**D**) Jumping and opening of the loop for *η* = 6 and *η* = 10 of SoftJM 3. (**E**) Curvature *κ* variation with aspect ratio *η* with normalized length (l/L) during SoftJM 3 bending and form closed loop. (**F**) Maximum jumping height (*H*_*max*_) per body length (BL) of SoftJM 3 at various *η* (n=3). (**G**) Maximum bending force (*F*_*bmax*_) to form a closed loop with aspect ratio *η* for SoftJM 3. (**H**) SoftJM 3 trajectories for different *η*. (**I**) Opening angle (*β*) as a function of time for different *η* for SoftJM 3.

To underscore the impact of aspect ratio *η* on jumping performance, we conducted systematic experiments on a range of *η* = 4 − 26 using SoftJM 3, with a fixed Young’s modulus (Fig. 6C). As expected, we find that the bending force 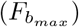 increases with a decrease in *η* (Fig. 6G), indicating increased energy storage at lower aspect ratios. Remarkably we observed that kinks also occur at *η*_*c*_*<*7 in SoftJM 3, similar to EPNs (Fig. 6C-F). Smaller *η* in SoftJMs resulted in quicker snapback (*η*=6, *β* opening from 0 to 120° in ∼5 ms) compared to higher *η* (*η* = 26, *β* opening from 0 to 120° in *>*100ms) (Fig. 6I and Movie S4).

After kink formation, the jumping height nearly doubled for *η* = 6 (∼ 16 BL) compared to *η* = 12 (pre-kink), suggesting that reversible kinks enhance jumping efficiency (Fig. 6F) by increasing the bending energy stored while ensuring a relatively reduced restoring force.

Thus, by actively modulating their aspect ratio to achieve a critical *η* for kink formation, EPNs store higher elastic energy and facilitate a latch between ventral-ventral contacts. Through corroborative experiments with SoftJM, we confirm that kinks at critical *η* lead to higher jumps and faster opening, while staying within a maximum force budget. This suggests that EPNs reach their upper force limits to bend and then kink to reduce force while continuing to store energy efficiently.

### Stiffness Improves Soft Jumper Performance

Our AFM measurements show that EPNs use stiffer structural materials, contributing to their superior jumping capabilities compared to *C. elegans* (Fig.3E). Recognizing the strategic importance of combining both softness and stiffness, we engineered a silicone-based SoftJM 4 by filling a 3D-printed mold with silicone and incorporating carbon fiber (CF), a much stiffer material (∼ 134 GPa), at its core to enhance the jumper’s stiffness (Fig.7A).

We conducted experiments with SoftJM 4, varying the number of CFs from 1-8, by bending the models into closed loops and releasing them (Fig.7B,F). We observe that take-off velocity (*V*_*max*_) increases linearly with the number of CFs used, reaching up to ∼ 13 m/s for CF 8, which also achieves the maximum jump height of (∼3 m) (Fig. 7B,C). The maximum power output (*P*_*max*_) reaches 20 W/Kg for CF8, which is 20x higher than the power generated by SoftJMs with just 1 CF (Fig. 7D). The addition of CFs minimally increases the mass but significantly enhances the strength of SoftJM, allowing it to store an increased bending force of approximately 7 N (Fig. 7E).

**FIG. 7.**
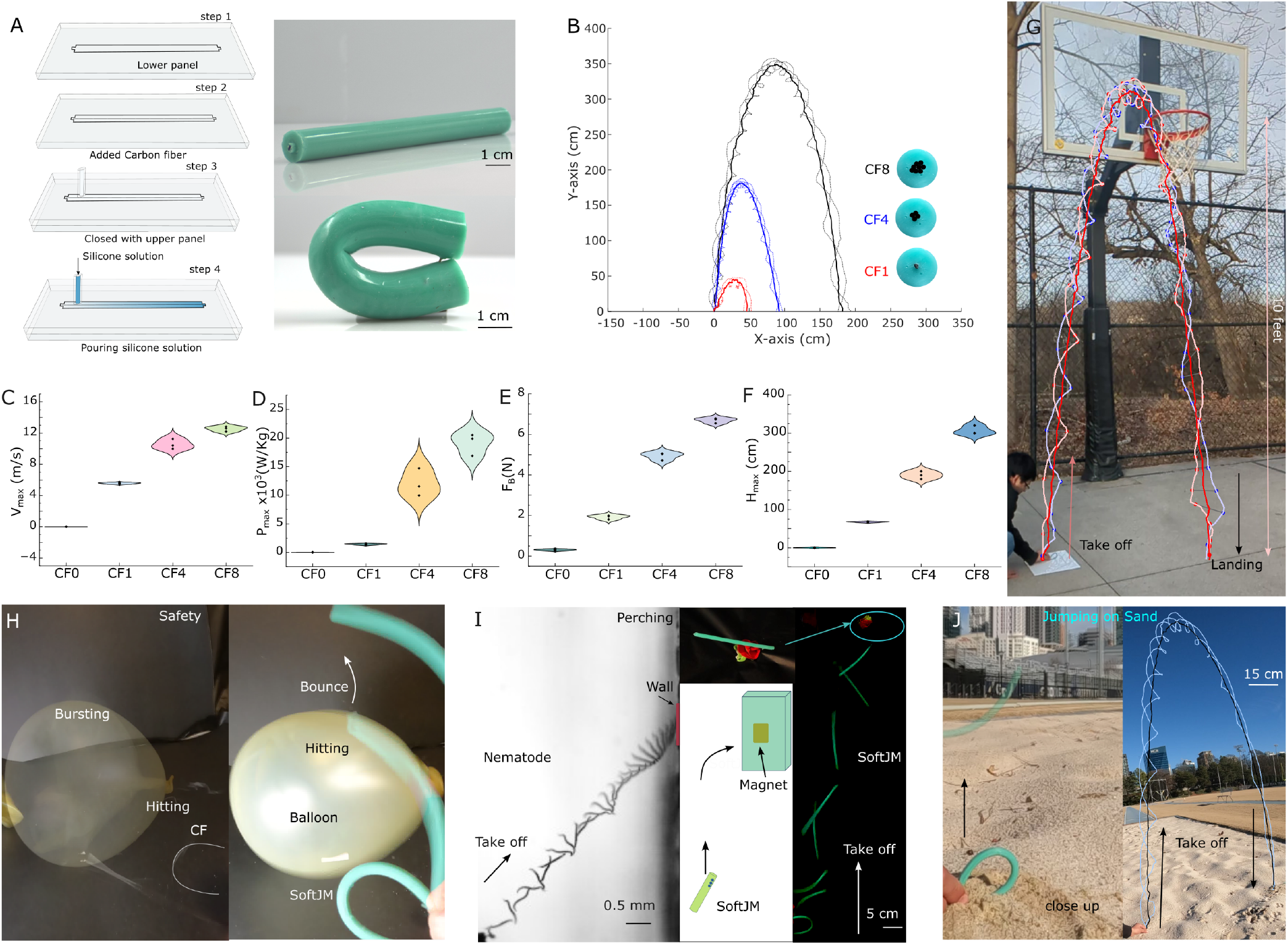
Versatile Applications and Safety Demonstrations of SoftJM 4. (**A**) Fabrication steps of SoftJM 4, showing the carbon fiber (CF) incorporated SoftJM, where *η* = 24 (details in Table S3). (**B**) Cross-sectional view of the SoftJM with 1, 4, and 8 carbon fibers and full trajectories of SoftJM 4 represented with red, blue, and black for CF1, CF4, and CF8, respectively, with highlighted lines showing centroid trajectories. (**C**) Maximum takeoff speed (*V*_*max*_) of SoftJM 4 with different numbers of CF (n=3). (**D**) Maximum Power (*P*_*max*_) generated by SoftJM 4 during takeoff. (**E**) Maximum bending force (*F*_*b*_) of SoftJM 4 (n=3). (**F**) Maximum height (*H*_*max*_) comparison of various soft jumping models. (**G**) SoftJM 4 (CF8) jumping the height of a basketball hoop (10-feet) . (**H**) Safety feature demonstrated by a carbon fiber opening up rapidly bursting a balloon, while the SoftJM 4 bounces on the balloon without damage. (**I**) Ambush foraging behavior of EPN towards hosts. Similarly, magnetic SoftJM 4 exhibits attraction towards a magnetic substrate. (**J**) SoftJM 4 can jump on granular substrates like sand.

### Versatility, Safety and Sand Jumping of SoftJM

We showcase the jumping capability of SoftJM 4 by demonstrating its ability to cross a ∼ 10-foot basketball net (Fig. 7G, Movie S5) and perform other tasks, such as jumping onto a car and hurdling (Fig. S7). Although the jumper packs a lot of power, it is safe around humans due to the strategic combination of softness and stiffness. We demonstrate that an air-filled balloon bursts upon being hit by just a naked CF rod (Fig. 7H, Movie S5). In contrast, when SoftJM 4 hits the balloon, it softly bounces back into the air, highlighting the added advantage of compliance in both jumping and safety.

Ambush foraging is a common strategy for EPNs to find hosts such as flying insects, as seen in EPNs attaching to surfaces after jumping (Fig. 7I, Movie S5). Similarly, we demonstrate perching behavior with a magnetic SoftJM, which is attracted to and attaches to a magnetic wall during a jump. We also show that the SoftJM can successfully perform jumping tasks on rough surfaces, such as sand, where it generates enough power to counter granular friction and leap, dispersing sand particles into the air (Fig. 7J, Movie S5). These multitasking features highlight the SoftJM’s flexibility and adaptability in different environments, underscoring its potential applications in soft robotics.

## Discussion

We investigate how the entomopathogenic nematode *Steinernema carpocapsae* exploits reversible kinks to enhance its aerial jumping capabilities and engineer bioinspired soft jumping robots. We validate our observations for varying center of mass through numerical simulations of a diverse initial worm configurations, complemented by systematic exploration of SoftJM performance as a function of mechanics, geometry and structural heterogeneity.

We estimate the spring forces generated in the curvature of EPN’s *α*-shaped body and the force of the liquid-latch holding the ventral-ventral contact. Our study of SoftJM reveals that reversible kink formation leads to a nonlinear spring response, resulting in a power output increase beyond what a similar actuation system in the linear bending regime would produce. Finally, we highlight the versatile jumping performance of various soft jumping models, situating both the EPN and SoftJM within the broader context of jumping organisms and robots.

### Bi-Directional Jumping in Flexible Rods

Jumping locomotion, common across various animals, typically utilizes limbs or specialized appendages. Examples include cats, humans, squirrels, fleas, springtails and spiders, with masses ranging from micrograms to tens of kilogram (Fig. 8) [33–36]. Limbless animals, such as worms, snakes, and larvae, adopt a closed-loop strategy, using their bodies as an ‘emergent limb’ for jumping [37–41]. Unlike limbed animals, these limbless creatures face geometric and muscular constraints that limit their ability to change their center of mass (COM) for jumping [32].

**FIG. 8.**
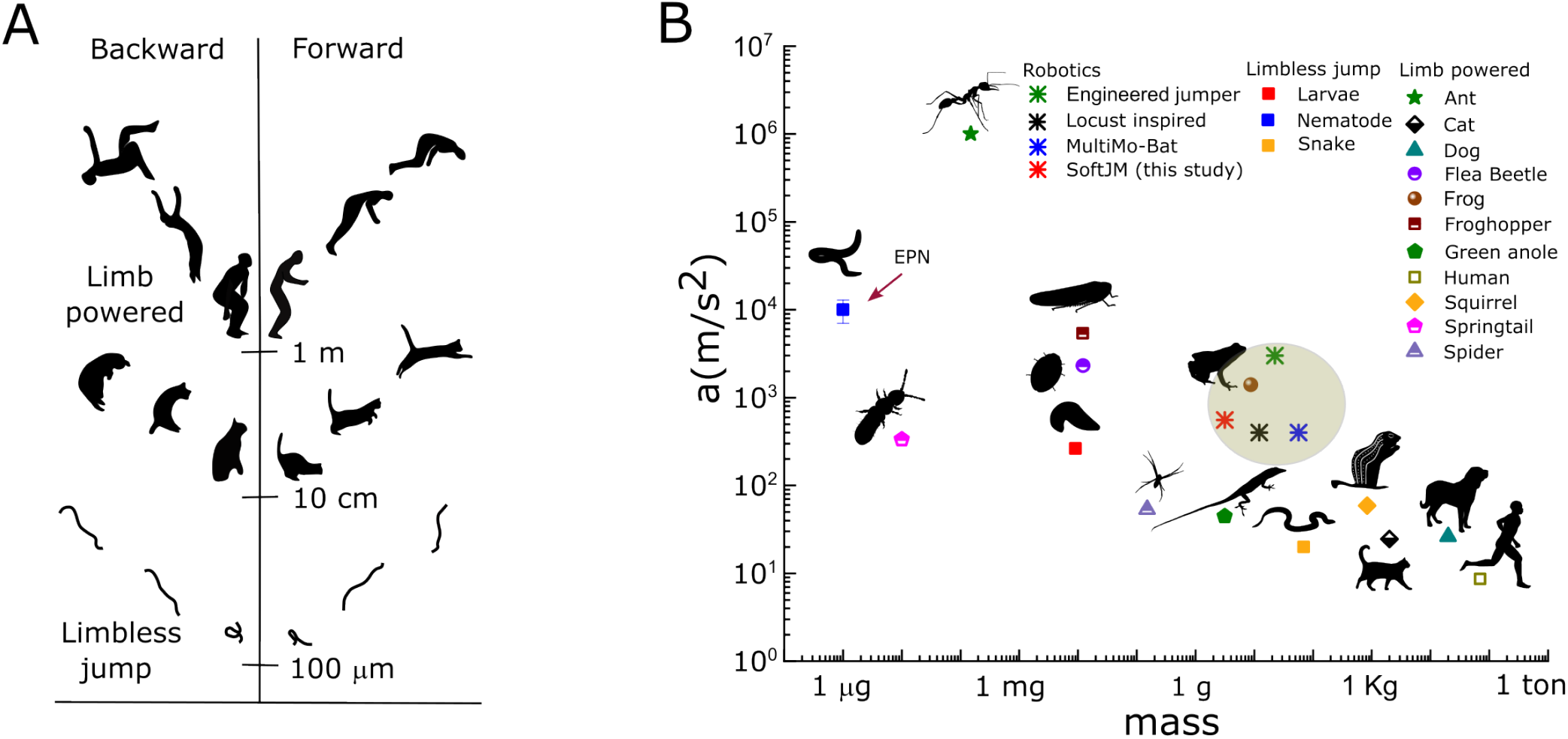
Bidirectional Jumping and Acceleration in Biological and Engineered Systems. (**A**) Forward and backward bidirectional jumping at various biological scales. (**B**) Maximum acceleration while jumping as a function of mass for different biological organisms and engineered models (Table S9).

We show that tiny, limbless nematodes can shift their COM using their *α*-shaped body configuration to jump both forward and backward, suggesting they might use such bi-directional control to locate suitable hosts (Fig.2A). The COM shifting in EPNs is governed by the head angle (*α*) and loop angle (*θ*), which together define the take-off angle (*γ*) and determine the jump’s direction.

EPNs can achieve bi-directional jumps with nearly equal heights of about (∼ 20 BL) (Fig. S1B). For context, humans can only achieve forward and backward jumps with heights *<* 1 BL (Fig.8A).

Building on this principle, we demonstrate that simple, limbless soft robots can also achieve bidirectional jumps. Our in-silico simulations reveal a rich phase space that serves as a design framework for flexible, curved, soft robot jumpers. This phase diagram opens new design modalities for limbless jumpers, enabling precise and controlled jumping in various directions using curvature and flexibility.

### Stiffer Cuticle Essential for Jumping

Nematode stiffness is key to facilitating low-power activities like swimming, crawling, and nictation, contrasting with the higher power demands of jumping [37]. Only a few nematode species can jump, raising questions about the structural stiffness fundamental for this feat and its role in storing muscle energy as a spring. We measured the stiffness of *C. elegans* and *S. carpocapsae* using AFM, finding that jumping EPNs exhibit greater stiffness compared to non-jumping *C. elegans* nematodes. The difference in nematode stiffness is further highlighted by comparing cuticle thickness to diameter ratios: *S. carpocapsae* has a ratio of 1:30, significantly thicker than *C. elegans* ratio of 1:88, indicating a much thicker cuticle in EPNs[25, 42] (Table S1). This cuticular thickening is pronounced during the dauer stage, the only life stage of the nematode that jumps - a trait essential to finding a host to complete its lifecycle. These structural adaptations, as our experiments with SoftJM suggest, provide a mechanical advantage by providing a stiff layer that supports the hydroskeleton, critical for storing energy in the body curvature necessary for jumping.

### SoftJM robots

Nature may not invent wheels, but it certainly evolves jumpers across taxa, from tiny sand fleas to giant humpback whales. Inspired by these agile systems, engineers have developed various jumping robots based on locusts, fleas, galagos, and others to navigate challenging terrains, including planetary surfaces [43–46]. However, these robots have faced challenges such as large sizes and masses, explosive power sources, safety concerns near humans and other living creatures, and multifunctional performance on complex surfaces such as granular matter [45, 47, 48].

Drawing inspiration from the jumping mechanics of limbless nematodes, we implemented two strategies to enhance jumping performance in our soft robotic structures. First, by exploiting kink instabilities, we doubled the jumping performance of our SoftJM 3 prototype, achieving up to ∼16 BL Fig. 6F). Second, by incorporating stiff carbon fiber rods into the soft silicone matrix, our SoftJM 4 prototypes (CF8, 3.5 g) increased the jumping height to 25 BL or 10 feet (basketball hoop height) (Fig.7).

Our proof-of-concept designs overcome previous limitations of jumping robots and introduce new design motifs. By using curvature for controlled directional take-offs, shielding sharp-pointed carbon fibers in the soft polymer for safe interactions, perching using small magnets and jumping on sandy substrates, we significantly expand the operational envelope of soft robots. The use of flexible rods as adaptive limbs opens possibilities for more sophisticated geometrical and topological instabilities that can be harnessed for new classes of adaptive jumping structures.

## Acknowledgements

We thank the Bhamla Lab for feedback. M.S.B. acknowledges funding support from NIH MIRA Grant R35GM142588; NSF Grants PHY-2310691; MCB-2313724; CMMI-2218382;CAREER iOS-1941933; and the Open Philanthropy Project. A.R.D acknowledges funding support from NIH MIRA Grant R35GM137934.

